# Autotaxin is a Potential Link between Genetic Risk Factors and Pathogenesis of Systematic Lupus Erythematosus in Plasmacytoid Dendritic Cells

**DOI:** 10.1101/2021.10.25.465592

**Authors:** Yumi Tsuchida, Hirofumi Shoda, Masahiro Nakano, Mineto Ota, Tomohisa Okamura, Kazuhiko Yamamoto, Makoto Kurano, Yutaka Yatomi, Keishi Fujio, Tetsuji Sawada

## Abstract

**Background:** The importance of autotaxin, an enzyme that catalyzes lysophospholipid production, has recently been recognized in various diseases, including cancer and autoimmune diseases. Herein, we examined the role of autotaxin in systemic lupus erythematosus (SLE), utilizing data from ImmuNexUT, a comprehensive database consisting of transcriptome data and expression quantitative trait locus (eQTL) data of immune cells from patients with immune-mediated disorders.

**Methods:** Serum autotaxin concentrations in patients with SLE and healthy controls (HCs) were compared. The transcriptome data of patients with SLE and age-and sex-matched HCs were obtained from ImmuNexUT. The expression of *ENPP2*, the gene encoding autotaxin, was examined in peripheral blood immune cells. Next, weighted gene correlation network analysis (WGCNA) was performed to identify genes with expression patterns similar to *ENPP2*. The ImmuNexUT eQTL database and public epigenomic databases were used to infer the relationship between autotaxin and pathogenesis of SLE.

**Results:** Autotaxin levels were elevated in the serum of patients with SLE compared to HCs. Furthermore, the expression of *ENPP2* was higher in plasmacytoid dendritic cells (pDCs) than in other immune cell subsets, and its expression was elevated in pDCs of patients with SLE compared to HCs. In WGCNA, *ENPP2* belonged to a module that correlated with disease activity. This module was enriched in interferon-associated genes and included genes whose expression was influenced by single-nucleotide polymorphisms associated with SLE, suggesting that it is a key module connecting genetic risk factors of SLE with disease pathogenesis. Analysis utilizing the ImmuNexUT eQTL database and public epigenomic databases suggested that the increased expression of *ENPP2* in pDCs from patients with SLE may be caused by increased expression of interferon-associated genes and increased binding of STAT3 complexes to the regulatory region of *ENPP2*.

**Conclusions:** Autotaxin may play a critical role in connecting genetic risk factors of SLE to disease pathogenesis in pDCs.

## Introduction

Lysophospholipids are phospholipids with only one fatty acid chain, such as sphingosine-1-phosphate, lysophosphatidic acid, and lysophosphatidylserine. These molecules play important roles in various biological processes by signaling through their respective G protein-coupled receptors^1^. Recently, enzymes that catalyze the formation of lysophospholipids, such as autotaxin and phosphatidylserine-specific phospholipase A1 (PS-PLA1), were reported as potential biomarkers of various diseases. For example, serum autotaxin levels and PS-PLA1 levels have been associated with melanoma^2^. They have also been associated with immune-mediated diseases; for example, their levels are elevated in the serum of patients with lupus nephritis compared to patients with other glomerular diseases^3^, and the level of PS-PLA1 in the serum correlated with disease activity in patients with systemic lupus erythematosus (SLE)^4^. Additionally, autotaxin is increased in the urine of patients with lupus nephritis^5^. Therefore, autotoxin may be important in the pathogenesis of SLE; however, its exact role in SLE remains unclear.

Both genetic and environmental factors contribute to the development of SLE. Various genome-wide association studies (GWAS) have been performed to identify genetic risk factors for SLE, and more than 100 loci have been reported to be associated with SLE^6^. However, the exact mechanism by which those GWAS SNPs contribute to the development of SLE in often not clear, and further studies are necessary to elucidate the link between genetic risk factors of SLE and disease pathogenesis and to develop new treatment strategies.

Recently, our group created a transcriptome and expression quantitative trait locus (eQTL) database of immune cells from various immune-mediated diseases, named “ImmuNexUT” ^7^. This database consists of the genome and transcriptome of 28 immune cells from patients with various immune-mediated diseases, as well as healthy controls (HCs). In this study, to further examine the role of autotaxin in SLE, we utilized data from ImmuNexUT^7^ to examine the expression pattern of autotaxin in immune cells, its relationship with clinical parameters and genetic risk factors of SLE.

## Methods

### Autotaxin ELISA

Serum samples were obtained from 54 patients with active, untreated SLE (42 females and 12 males) who fulfilled the 1997 American College of Rheumatology revised classification criteria at the University of Tokyo Hospital and Tokyo Medical University Hospital. Serum samples were also collected from 237 healthy individuals (69 females and 168 males).

Serum autotaxin levels were quantified using a one-step immunoenzymatic assay, as described previously^8^. Briefly, rat monoclonal antibodies specific for human autotaxin were used in the two-site immunoassay. Magnetic polymer beads coated with the primary monoclonal antibody and an alkaline phosphatase-labeled secondary monoclonal antibody were placed in the reaction cup. After adding the diluted serum sample to the reaction cup, the sample was evaluated using a commercial automated immunoassay analyzer (AIA-2000 system; Tosoh, Tokyo, Japan).

### Differential expression analysis

For the differential gene expression analysis, RNA-seq data from the ImmuNexUT study^7^ were utilized. Details of sample collection, RNA sequencing, and data processing have been described in detail previously^7^. Patients with SLE with expression data for plasmacytoid dendritic cells (pDCs) after filtering (n = 59) and a subset of HCs, age- and sex-matched to the patients with SLE (n = 51) were selected. Patients with SLE fulfilled the American College of Rheumatology 1997 classification criteria^9^ and SLICC 2012 criteria^10^. Patients with malignancies, acute infections, or those taking more than 20 mg of prednisolone were excluded from the study. Disease activity was assessed using SLEDAI-2K^11^. Differential expression analysis was performed using edgeR after trimming the mean of M-values (TMM) normalization^12^.

### WGCNA

For weighted gene co-expression network analysis (WGCNA), genes with read counts of less than 10 in 90% or more of the samples were excluded. The count data were normalized with TMM normalization and converted to log 2 (CPM + 1). WGCNA was performed using WGCNA package version 1.68 with signed network and soft thresholding power set to 8. One sample was excluded as an outlier. The representative expression value of each module, “eigengenes,” was calculated, and the correlation between these eigengenes and various clinical parameters was assessed. Pathway analysis was performed using DAVID^13^ with the REACTOME pathways^14^.

### eQTL analysis

For eQTL analysis, eQTL data from the ImmNexUT database were utilized, and the details are described in our previous study^7^. After filtering out genes with low expression in each cell subset, the expression data were normalized using TMM normalization, converted to CPM, and normalized across samples using an inverse normal transform. Probabilistic estimation of expression residuals (PEER)^15^ was used to detect hidden covariates. A QTLtools permutation pass with 10,000 permutations was used to obtain gene-level nominal P value thresholds corresponding to a false discovery rate < 0.05 for each cell subset. Next, forward-backward stepwise regression eQTL analysis was performed with a QTLtools conditional pass. ATAC-seq data provided by Calderon et al.^16^ were used to assess the relationship between eQTL SNPs and ATAC-seq peaks. Candidate cis-regulatory elements from the ENCODE study were used to visualize distal enhancer-like signatures^17^, and data from JASPAR^18^ were used to visualize predicted transcription factor binding sites.

### Ethics declarations

This study was performed in accordance with the principles of the Declaration of Helsinki. The ethics committees at the University of Tokyo Hospital and Tokyo Medical University Hospital approved this study, and written consent was obtained from all participants (G10095).

### Statistics

Categorical data were analyzed using Fisher’s exact test. Differences between two groups were compared using Student’s *t*-tests for normally distributed continuous data and Mann–Whitney U tests for non-normally distributed continuous data. Correlations were evaluated using Spearman’s rank correlation coefficients. Statistical significance was set at P < 0.05. Statistical analyses were performed using R version 3.5.0 (R Foundation for Statistical Computing).

## Results

### Autotaxin is elevated in the serum of patients with SLE

We examined the concentration of autotaxin in the serum of untreated patients with SLE and in the serum of HCs. As the serum concentration of autotaxin is reported to be different between females and males^8^, data from each sex group were analyzed separately. The serum concentration of autotaxin was higher in patients with SLE than in HCs in both females (Figure 1a) and males (Figure 1b).

**Figure 1.**
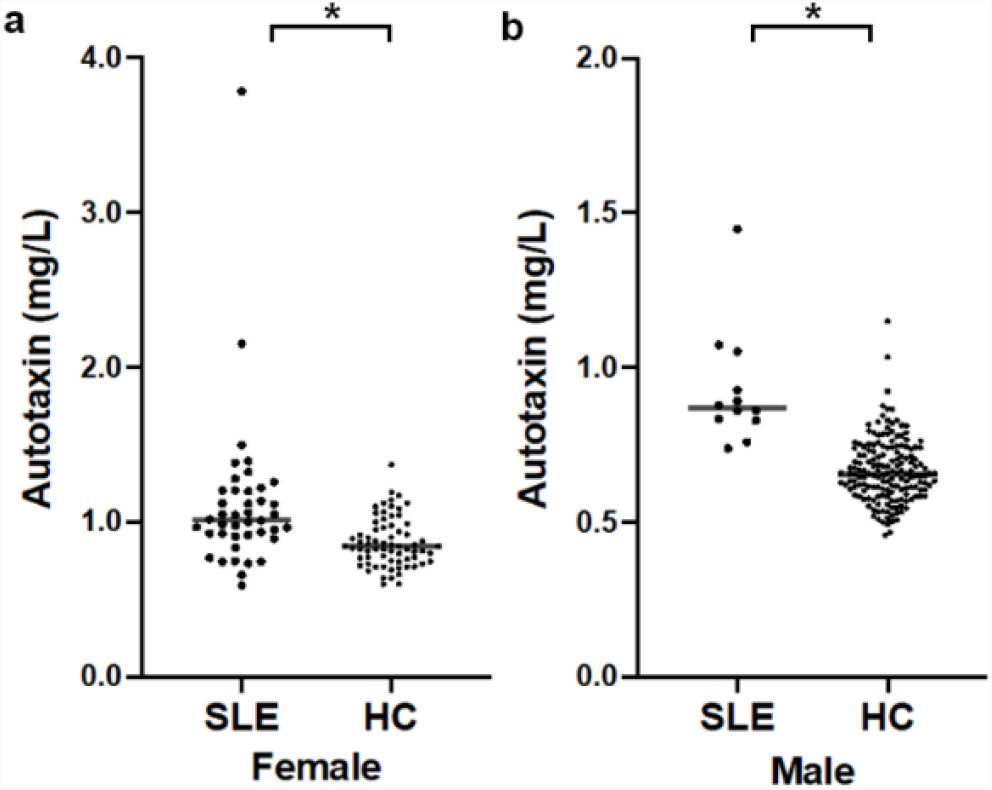
Autotaxin is elevated in the serum of patients with SLE. Serum autotaxin levels in (a) female and (b) male patients with SLE and HCs.

### The expression of ENPP2 is elevated in plasmacytoid dendritic cells of patients with SLE

Next, we utilized the transcriptome data of patients with SLE and age- and sex-matched HCs from ImmuNexUT to examine the mRNA expression level of *ENPP2*, which encodes autotaxin. The transcript levels of *ENPP2* were higher in pDCs compared to other immune cell subsets, as reported previously^7, 19^ (Figure 2a). Therefore, we focused on pDCs. The expression of *ENPP2* was elevated in pDCs of patients with SLE compared to HCs, as noted by others^19^ (Figure 2b). When the transcriptome of pDCs from females and males were analyzed separately, *ENPP2* was differentially expressed in both sexes (Figure 2c and 2d).

**Figure 2.**
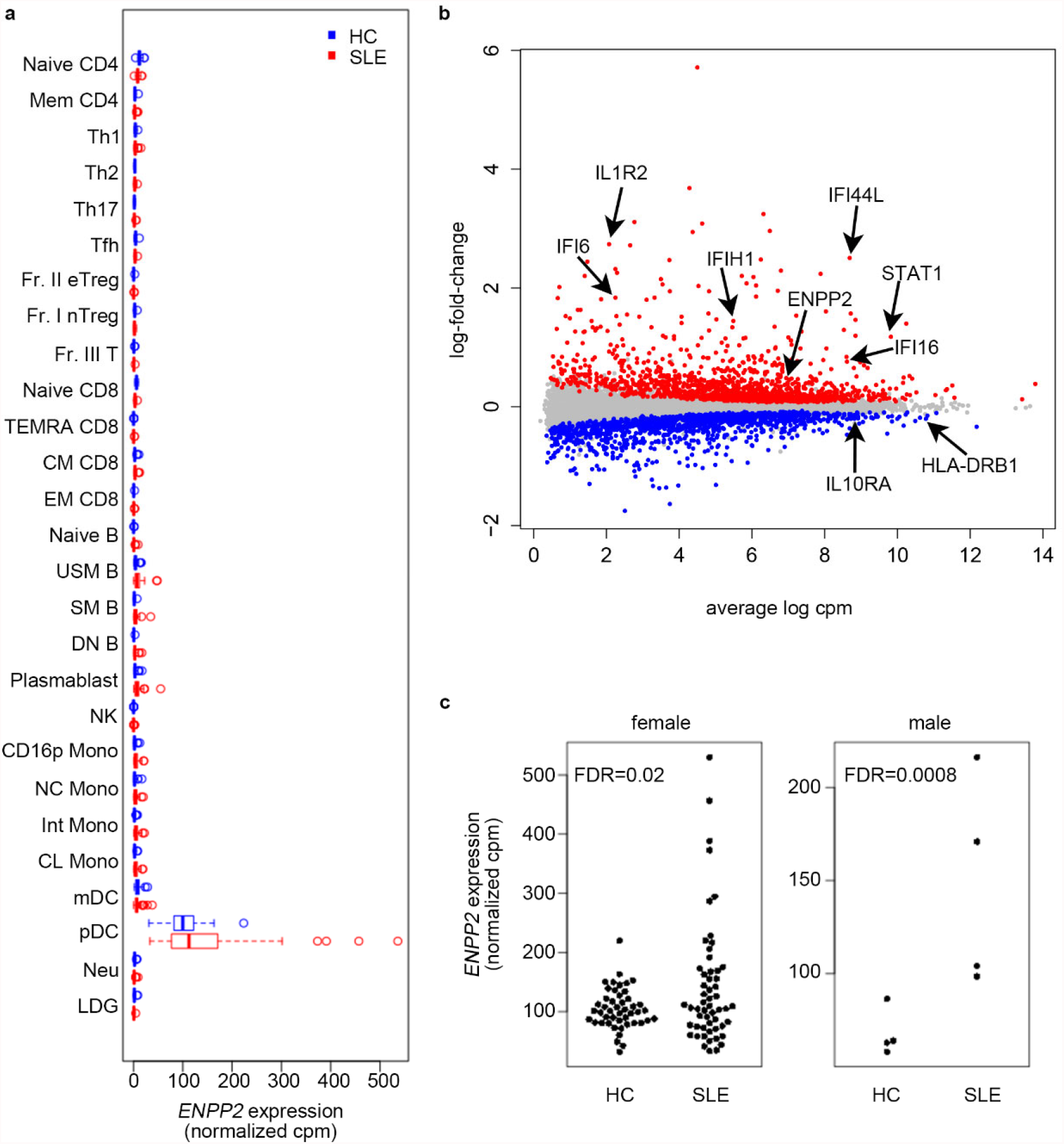
The expression of ENPP2 is elevated in plasmacytoid dendritic cells of patients with SLE. (a) Expression of *ENPP2* in immune cell subsets of patients with SLE and HCs. For abbreviations, see Supplementary Table S1. (b) MA plot of pDCs from patients with SLE and HCs with selected genes highlighted. (c) *ENPP2* expression in pDCs from HCs and patients with SLE.

### ENPP2 is co-expressed with interferon-associated genes

To further examine the role of *ENPP2* in the pathogenesis of SLE, WGCNA was performed to group genes into modules of genes with similar expression patterns, using transcriptome data of pDCs from patients with SLE (Figure 3a). *ENPP2* belonged to the magenta module (Supplementary Table S2), the eigengene score of which was correlated with SLEDAI-2K and with anti-RNP antibody positivity (Figure 3a). The eigengene score of this module was higher in patients with moderate or high disease activity (SLEDAI-2K > 6) compared to patients with mild or no disease activity (SLEDAI-2K ≤ 6) (Figure 3b). The magenta module included many interferon-associated genes, such as *IFIH1* and *ISG15*; pathway analysis of the genes in this module showed enrichment of interferon-associated pathways (Figure 3c).

**Figure 3.**
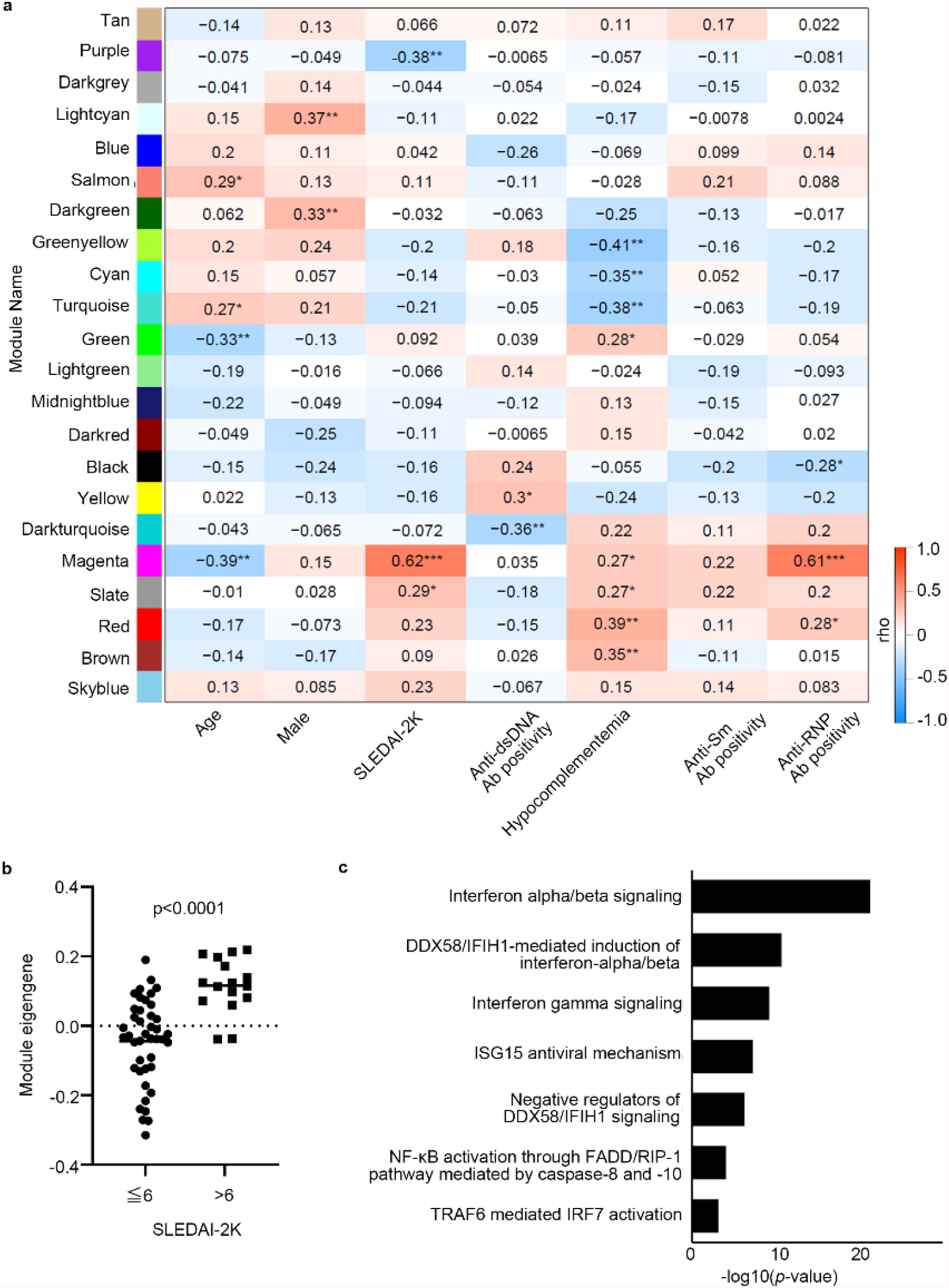
*ENPP2* is co-expressed with interferon-associated genes. (a) Correlation of WGCNA modules with clinical parameters. The numbers indicate the correlation coefficients. * indicates p < 0.05, ** indicates p < 0.01, and *** indicates p < 0.001. (b) Module eigengenes of the magenta module in patients with SLE with SLEDAI-2K scores ≤6 and >6. (c) Pathway analysis of the members of the magenta module using REACTOME pathways. Data were analyzed using Fisher’s exact test.

### The magenta module links genetic risk factors of SLE with the pathogenesis of SLE

Further examination of the members of the magenta module using the eQTL database from ImmuNexUT^7^ indicated that this module contains genes whose expression is influenced by SLE GWAS SNPs in pDCs. For example, rs13385731, in linkage disequilibrium (LD) with SLE GWAS SNP rs13425999^20^, is an eQTL of *RASGRP3* in pDCs (Figure 4a). In other words, in patients with the SLE risk allele, the expression of *RASGRP3* is increased in pDCs. In addition, rs11059921, in LD with SLE GWAS SNP rs11059919^21^, influences the expression of *SLC15A4*, suggesting that this module may reflect a link between genetic risk factors for SLE and its disease pathogenesis (Figure 4b).

**Figure 4.**
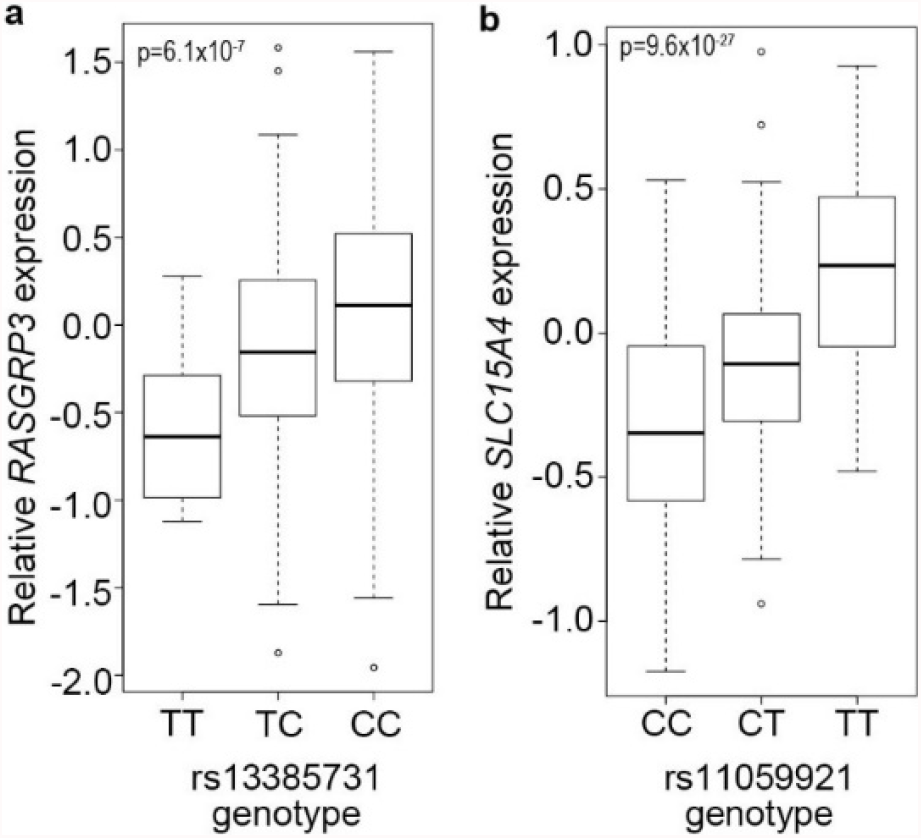
The magenta module links genetic risk factors of SLE with the pathogenesis of SLE. (a) eQTL effect of rs13385731 on *RASGRP3*. (b) eQTL effect of rs11059921 on *SLC15A4*.

### rs11778951 influences the expression of ENPP2 in pDCs

Next, to elucidate the mechanism underlying the increased expression of *ENPP2* in pDC of patients with SLE, we examined the eQTL database in ImmuNexUT. rs11778951, located in the *ENPP2* gene, influenced the expression of *ENPP2* in pDCs with the G allele increasing the expression of *ENPP2* (Figure 5a). ATAC-seq data of immune cells from a public database^16^ indicated that rs11778951 coincides with a peak specific to pDCs (Figure 5b). This locus was predicted to be a distal enhancer-like signature in ENCODE^22^ and resides near predicted IRF1 and STAT3 binding sites^18^. Furthermore, rs11778951 alters a STAT-binding motif at this site^23^. Therefore, the increase in interferon signaling in patients with SLE may cause increased binding of STAT3 complexes to this site in pDCs, leading to increased expression of *ENPP2*.

**Figure 5.**
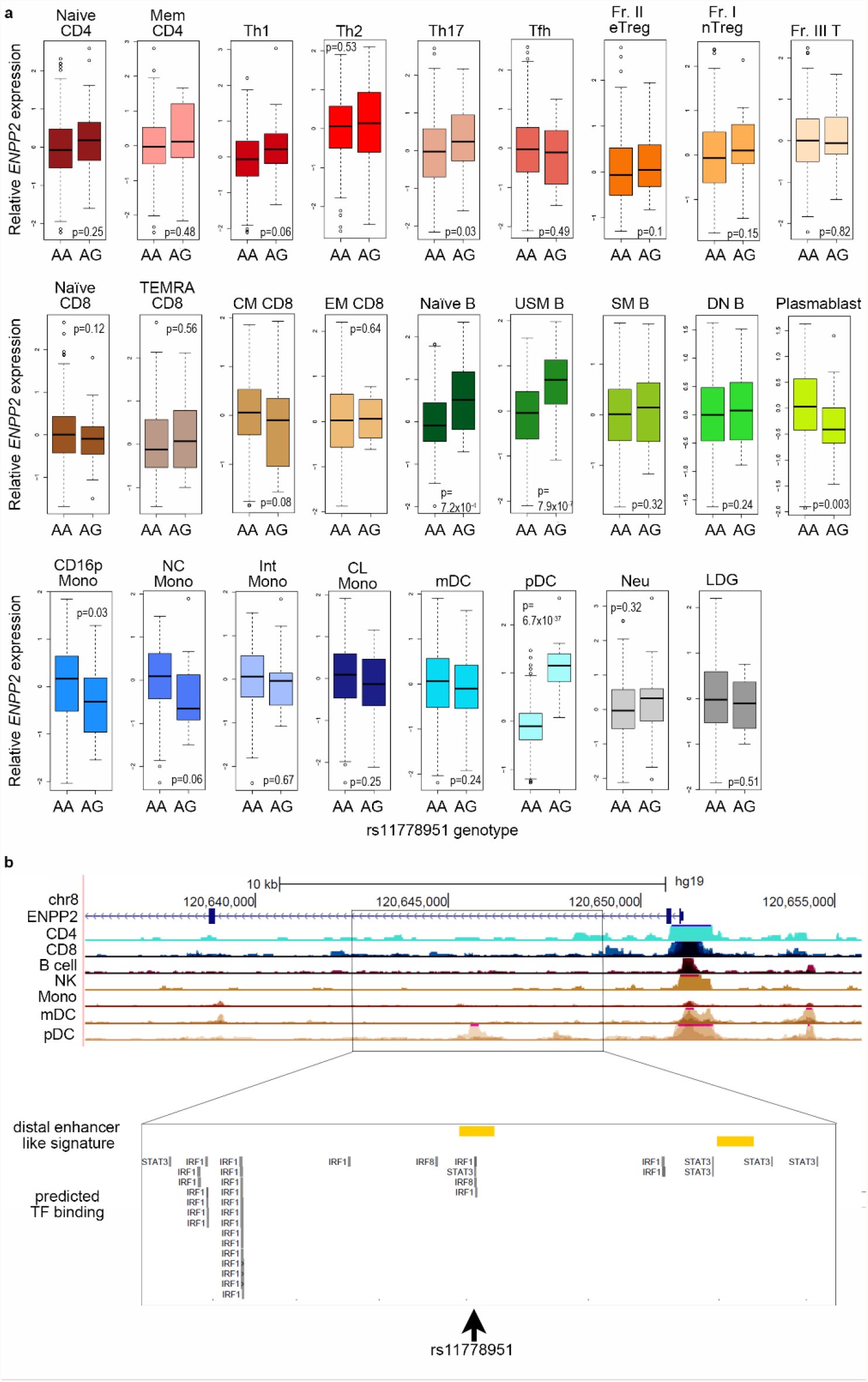
rs11778951 influences the expression of *ENPP2* in pDCs. (a) Expression of *ENPP2* by rs11778951 genotype. (b) ATAC-seq peaks near rs11778951 are shown, along with distal enhancer-like signatures and predicted transcription factor (TF) binding sites.

## Discussion

We analyzed autotaxin and related pathways using the ImmuNexUT database in association with SLE. The level of autotaxin was elevated in the serum of patients with SLE, and the expression of *ENPP2*, encoding autotaxin, was increased in pDCs of patients with SLE. In WGCNA, *ENPP2* belonged to the magenta module, whose members are enriched in interferon-associated genes and whose expression correlates with disease activity of SLE. The magenta module included genes whose expression was affected by SLE GWAS SNPs, suggesting that *ENPP2* may be one of the key molecules linking genetic risk factors of SLE with disease pathogenesis. Furthermore, rs11778951, which has an eQTL effect on *ENPP2*, coincides with an ATAC-seq peak specific to pDCs and alters STAT protein binding motifs, suggesting that the increased interferon signaling in patients with SLE may contribute to the increased expression of *ENPP2* in SLE.

Our results suggest that autotaxin plays a role the pathogenesis of SLE. It has been reported that autotaxin is highly expressed in the adipose tissue, central nervous system, and reproductive organs, and in the immune system, autotaxin is expressed by endothelial cells in high endothelial venules^24^. Although pDCs have not generally gained much attention as an important source of autotaxin, a recent report on the role of autotaxin in COVID-19 suggested that autotaxin expressed by pDCs affects the development and function of pDCs by regulating the local production of lysophosphatidic acid^19^. Our study suggests that the increased expression of autotaxin in pDCs of patients with SLE may contribute to the pathophysiology of SLE by altering pDC function as well.

Our study indicated that the expression of interferon-associated genes show correlation with the expression of *ENPP2* in pDCs. Our results are consistent with those of a previous study demonstrating that type I interferons are involved in the production of autotaxin^25^. Furthermore, by combining WGCNA with the eQTL database, a potential mechanism for the increased expression of *ENPP2* in pDCs in patients with SLE was identified. That is, the increase in interferon-signaling induced by genetic risk factor of SLE may lead to the increased expression of *ENPP2* in pDCs by increasing the binding of STAT proteins to rs11778951, an eQTL locus of *ENPP2* in pDCs.

One major limitation of this study is that the expression of autotaxin in pDCs was only assessed at the mRNA level. Further studies examining the expression of autotaxin at the protein level and its function in pDCs would further clarify the role of autotaxin in the pathogenesis of SLE and provide insights for identifying therapeutic targets.

In conclusion, our transcriptome analysis of pDCs identified a gene module whose expression correlates with disease activity in patients with SLE and is affected by SLE GWAS SNPs. By examining genes in this module, autotaxin expressed by pDCs was identified as a possible key molecule in the link between genetic risk factors and the pathogenesis of SLE.

## Acknowledgements

We would like to express our gratitude to all the study participants for their cooperation for this study. The supercomputing resource, SHIROKANE, was provided by the Human Genome Center, The University of Tokyo.

## Data availability

The RNA-seq data is available at the National Bioscience Database Center (NBDC) (E-GEAD-397).

