## Supplementary Tables for "Autotaxin is a Potential Link between Genetic Risk Factors and Pathogenesis of Systematic Lupus Erythematosus in Plasmacytoid Dendritic Cells"

### Supplementary Table S1. Cell subset definitions

| Subset name | Abbreviation | Definition |
|---|---|---|
| **CD4 T cells** | | |
| Naïve CD4 T cells | Naïve CD4 | CD3+/CD4+CD8-/CCR7+CD45RA+ |
| Memory CD4 T cells | Mem CD4 | CD3+/CD4+CD8-/non-naive CD4+/CD25- |
| T helper 1 cells | Th1 | CD3+/CD4+CD8-/non-naive CD4+/CD25-/CXCR5-CCR6-/CXCR3+CCR4- |
| T helper 2 cells | Th2 | CD3+/CD4+CD8-/non-naive CD4+/CD25-/CXCR5-CCR6-/CXCR3-CCR4+ |
| T helper 17 cells | Th17 | CD3+/CD4+CD8-/non-naive CD4+/CD25-/CXCR5-CCR6+/CXCR3- |
| T follicular helper cells | Tfh | CD3+/CD4+CD8-/non-naive CD4+/CD25-/CXCR5+ |
| Fraction II effector regulatory T cells | Fr. II eTreg | CD3+/CD4+CD8-/CD25++CD45RA- |
| Fraction I naïve regulatory T cells | Fr. I nTreg | CD3+/CD4+CD8-/CD25+CD45RA+ |
| Fraction III non-regulatory T cells | Fr. III T | CD3+/CD4+CD8-/CD25+CD45RA- |
| **CD8 T cells** | | |
| Naïve CD8 T cells | Naïve CD8 | CD3+CD19-/CD4-CD8+/CD45RA+CCR7+ |
| CD8+ T effector memory CD45RA+ cells | TEMRA CD8 | CD3+CD19-/CD4-CD8+/CD45RA+CCR7- |
| Effector Memory CD8 T cells | EM CD8 | CD3+CD19-/CD4-CD8+/CD45RA-CCR7- |
| Central Memory CD8 T cells | CM CD8 | CD3+CD19-/CD4-CD8+/CD45RA-CCR7+ |
| **B cells** | | |
| Naïve B cells | Naïve B | CD3-CD19+/IgD+CD27- |
| Unswitched memory B cells | USM B | CD3-CD19+/IgD+CD27+ |
| Switched memory B cells | SM B | CD3-CD19+/IgD-CD27+/CD38- |
| Double Negative B cells | DN B | CD3-CD19+/IgD-CD27- |
| Plasmablasts | Plasmablast | CD3-CD19+/IgD-CD27++/CD38+ |
| **Natural Killer cells** | NK | CD3-CD19-/CD14-/CD56+ |
| **Monocytes** | | |
| CD16 positive monocytes | CD16p Mono | CD3-CD19-/HLARDR+/CD56-/CD14+CD16+ |
| Non-classical monocytes | NC Mono | CD3-CD19-/HLARDR+/CD56-/CD14dimCD16+ |
| Intermediate monocytes | Int Mono | CD3-CD19-/HLARDR+/CD56-/CD14++CD16+ |
| Classical monocytes | CL Mono | CD3-CD19-/HLARDR+/CD56-/CD14+CD16- |
| **Dendritic cells** | | |
| Myeloid dendritic cells | mDC | CD3-CD19-/HLARDR+/CD56-/CD14-CD16-/CD11c+CD123- |
| Plasmacytoid dendritic cells | pDC | CD3-CD19-/HLARDR+/CD56-/CD14-CD16-/CD11c-CD123+ |
| **Neutrophils** | Neu | Immune-magnetically sorting with the "MACSxpress Neutrophil isolation Kit, human" |
| Low-Density Granulocytes | LDG | SSChigh /CD16+CD15+/CD14- |

**Supplementary Table S2. Members of the magenta module**

| | | | | | |
|---|---|---|---|---|---|
| RSAD2 | CMPK2 | USP18 | IFIH1 | TNFSF10 | PLSCR1 |
| SAMD9L | HESX1 | EPSTI1 | OASL | PARP12 | IFITM1 |
| ISG15 | OAS2 | SPATS2L | LAP3 | IFI16 | UBE2L6 |
| PARP9 | MX2 | PARP14 | SP100 | STAT1 | NMI |
| HERC6 | OAS1 | MNDA | HERC5 | MX1 | LY6E |
| GMPR | IFI27 | LGALS9 | ARHGEF3 | IFI44 | TRANK1 |
| LGALS3BP | IFIT1 | EIF2AK2 | IFIT3 | HSH2D | C19orf66 |
| COX5A | CD38 | CHMP5 | LOC100419583 | FBXO6 | CXCR2P1 |
| DTX3L | STAP1 | OAS3 | DDX60L | SP140 | DDX60 |
| PRKAG2 | SP110 | SLC38A5 | IFI35 | MAP2K6 | TRAFD1 |
| TAP1 | NT5C3A | SAMD9 | TNFSF13B | TMCC3 | ADAR |
| NAPA | NAMPT | MYD88 | DHX58 | IFI6 | IFITM2 |
| FIG4 | XAF1 | PDE7A | IFI44L | HLA.E | C5orf56 |
| PSMA2 | TRIM21 | BST2 | IFIT5 | PHF11 | IGSF6 |
| TRIM22 | LGMN | RUFY4 | FAM46A | IFITM3 | SCLT1 |
| TDRD7 | RGL1 | PIM3 | NEXN | SLFN12 | PI4K2B |
| PANK2 | SHISA5 | PPA1 | WDFY1 | SNX6 | NRIR |
| PNPT1 | RAB29 | TGM1 | NDUFA9 | BCL2L13 | TRIM5 |
| RASGRP3 | EPB41L3 | PML | PFKP | B2M | CKS2 |
| DDX58 | VSIG10L | CORO2B | SMC6 | ATP13A1 | INTS12 |
| TYMP | RTP4 | DYNLT1 | BLVRA | FYB | ZNFX1 |
| MASTL | KCTD19 | IFIT2 | BISPR | IRF2 | LOC101927027 |
| RUNX3 | ODF3B | SYNGR2 | GCH1 | LOC100505549 | EHD4 |
| EXOSC9 | SP140L | STAT2 | AP1M1 | DNAH12 | SUB1 |
| MESDC2 | P2RY6 | ARID5A | HOXB2 | CHST12 | PPIF |
| ZC3H12D | IGFBP4 | RCN1 | CPEB2 | SLC15A4 | BRCA2 |
| HIVEP3 | ICAM3 | GPR180 | TFEC | LAMP3 | LOC101927043 |
| RBCK1 | SSRP1 | NFKBIE | LAG3 | C3AR1 | SNX20 |
| JADE2 | UAP1 | RELB | PMAIP1 | PDCD2L | SPN |
| FCHSD2 | MRPL32 | BTG1 | DNAJC9 | C4orf33 | CASP10 |
| ISG20 | MCOLN2 | LINC00487 | CASP7 | CNP | JUP |
| TREX1 | CD69 | GALNT2 | HLA.F | AGRN | VRK3 |
| CHRNA1 | STAP2 | PARP1 | APOBEC3G | NUSAP1 | GEMIN2 |
| CD180 | MFN1 | SLC35A4 | SLC31A2 | HLA.A | PHACTR2 |
| ENPP2 | STAT4 | RBM43 | STX17 | RHOH | PIGB |
| NOC3L | CD274 | STRADA | GLRX | KLHDC7B | S100A3 |
| HNRNPD | RALY | C14orf1 | NFE2L3 | UBA7 | COQ2 |
| NRSN2.AS1 | BRIP1 | CALR | GRPEL1 | CD83 | MRPL44 |
| CHRNB1 | TMED3 | CARS | PAK1 | DCBLD1 | EIF3M |
| LRRC3 | TCN2 | TOLLIP | GTPBP1 | LTA | FN3KRP |
| MTX2 | MOV10 | TOR3A | TTC38 | SEMA4A | CAST |

| | | | | | |
|---|---|---|---|---|---|
| CBWD2 | APOL6 | IL2RG | GLT1D1 | PARP8 | C18orf8 |
| TRIM25 | NIPSNAP1 | PATL2 | GABARAPL1 | HTATIP2 | UBC |
| MICB | LMO2 | ICAM1 | NME8 | NRBP1 | TM2D2 |
| HEXA | SLC20A1 | CSTF2 | LDLRAP1 | PARP15 | PNO1 |
| SLC25A17 | SLC25A25 | RNF135 | CASP1 | ANKIB1 | SCML4 |
| SMARCD2 | TNS1 | CASP4 | TAF1B | GCNT2 | GBA |
| APOL1 | PI4KB | CHAC2 | NCF2 | ROBO1 | ITPR1 |
| SCPEP1 | NOP2 | TOR1B | TNF | STK40 | HCP5 |
| COBL | JHDM1D.AS1 | RBBP6 | TPX2 | WFDC21P | CCR7 |
| CDC42EP2 | CDC40 | DCAF11 | FAS | TGIF2 | STX11 |
| FAM13A | PTPN2 | PPP2R2A | CCDC62 | CELSR1 | TNFRSF10B |
| TMEM156 | SMAD1 | CENPE | DYNC2LI1 | TMEM255A | MILR1 |
| ASRGL1 | BIRC3 | DEK | LOXL4 | LRRCC1 | ANXA5 |
| CNKSR3 | GBP1 | TMEM17 | OTULIN | ELL2 | TOX4 |
| SLC44A4 | MKI67 | HIPK2 | MB21D1 | TRIM69 | UVRAG |
| PARP11 | CD48 | ZCCHC2 | PLVAP | MVK | IKZF3 |
| PRR11 | KMO | FRRS1 | LOC101927272 | CHD1L | OPTN |
| MST1R | MAX | MLLT3 | GBP4 | HDX | S100A5 |
| COG4 | L3MBTL4 | PLSCR4 | WHSC1L1 | SLC9A1 | P2RX5.TAX1BP3 |
| CNR2 | OR52N4 | NEDD4L | BTN2A2 | MED28 | MYO7B |
| HLA.F.AS1 | KCNS3 | HLA.DOB | B3GNT7 | E2F3 | HIST1H3H |
